# Environmental genomics points to non-diazotrophic *Trichodesmium* species abundant and widespread in the open ocean

**DOI:** 10.1101/2021.03.24.436785

**Authors:** Tom O. Delmont

## Abstract

Filamentous and colony-forming cells within the cyanobacterial genus *Trichodesmium* might account for nearly half of nitrogen fixation in the sunlit ocean, a critical mechanism that sustains plankton’s primary productivity at large-scale. Here, we report the genome-resolved metagenomic characterization of two newly identified marine species we tentatively named ‘Ca. *Trichodesmium miru*’ and ‘Ca. *Trichodesmium nobis*’. Near-complete environmental genomes for those closely related candidate species revealed unexpected functional features including a lack of the entire nitrogen fixation gene apparatus and hydrogen recycling genes concomitant with the enrichment of nitrogen assimilation genes and apparent acquisition of the *nirb* gene from a non-cyanobacterial lineage. These comparative genomic insights were cross-validated by complementary metagenomic investigations. Our results contrast with the current paradigm that *Trichodesmium* species are necessarily capable of nitrogen fixation. The black queen hypothesis could explain gene loss linked to nitrogen fixation among *Trichodesmium* species, possibly triggered by gene acquisitions from the colony epibionts. Critically, the candidate species are not only widespread in the 3-2000 μm planktonic size fraction of the surface of the oceans and seas, but might also substantially expand the ecological niche of *Trichodesmium*, stressing the need to disconnect taxonomic signal for this genus from a microbial community’s ability to fix nitrogen. Especially, differentiating diazotrophic from non-diazotrophic populations when counting *Trichodesmium* filaments and colonies might help refine our understanding of the marine nitrogen balance. While culture representatives are needed to move beyond metagenomic insights, we are reminded that established links between taxonomic lineages and functional traits might not always hold true.

## Introduction

Plankton in the sunlit ocean includes a wide range of microbial lineages with different functional capabilities that influence global biogeochemical cycles and climate^1–6^. The primary productivity of plankton is constrained by the amount of bioavailable nitrogen^7,8^, a critical element for cellular growth and division. Few bacterial and archaeal populations can code for the catalytic (*nifHDK*) and biosynthetic (*nifENB*) proteins required for biological nitrogen fixation, transferring a valuable source of nitrogen from the atmosphere to the plankton^9–11^. These populations are called diazotrophs and represent key marine players that sustain plankton primary productivity in large oceanic regions^9^.

Cyanobacterial species within the genus *Trichodesmium* first described in 1830^12^ are among the most prominent marine nitrogen fixers^13^, possibly accounting for half the biological nitrogen fixation in the sunlit ocean^14–16^. While other makers provide different trends (e.g., in^17^), phylogeny of the 16S rRNA genes points to three distinct *Trichodesmium* clades covering *T. thiebautii* and *T. hildebrandtii* (clade I), *T. tenue* and *T. contortum* (clade II), and *T. erythraeum* and *T. havanum* (clade III)^18^. For now, only cultures of *T. erythraeum* and *T. thiebautti* have been characterized with genomics. Insights from culture representatives and oceanic expeditions have provided a wealth of information regarding their biogeography, functional life styles and nitrogen fixation regulation mechanisms^14,19^. Most notably, *Trichodesmium* cells are capable of forming large blooms of filaments and colonies in the sunlit ocean^13,20–22^, regulate nitrogen fixation rates on a daily basis using a dedicated circadian rhythm^23,24^ as well as an associated microbiome^25–27^, and possess genes linked to the recycling of hydrogen, a by-product of nitrogen fixation^28–30^. Decades of scientific insights have shaped a dogma depicting the numerous *Trichodesmium* cells as carbon and nitrogen fixers with a highly beneficial role for plankton productivity.

Recent large-scale single cell and genome-resolved metagenomic surveys have dramatically expanded the genomic characterization of free-living marine microbes^31–34^ and lead to new insights into primary processes in the surface of the open ocean, including nitrogen fixation^34^. However, a major focus on small planktonic size fractions typically excluded organisms such as *Trichodesmium* that form filaments and colonies and occur in larger size fractions^21^. This gap is partially filled by a recent metagenomic survey that reconstructed and manually curated more than one thousand bacterial genomes abundant in large size fractions of the *Tara* Oceans expeditions^35^, which included five near-complete environmental *Trichodesmium* genomes. To our knowledge, they correspond to the first *Trichodesmium* genomes recovered without the need for cultivation, providing a new venue to study the ecology and evolution of populations within this genus and occurring in the open ocean. Here, we show that in addition to three genomes that resolve to *T. erythraeum* and *T. thiebautti*, this collection reveals two widespread *Trichodesmium* species most closely related to *T. tenue* and *T. contortum* (clade II) that display functional repertoires denoting a lack of the key ability fixing nitrogen.

## Results

### A first set of Trichodesmium environmental genomes from the sunlit ocean

A genome-resolved metagenomic survey was recently performed to target planktonic populations abundant in polar, temperate, and tropical sunlit oceans using nearly one thousand metagenomes (total of 280 billion reads) derived from the *Tara* Oceans expeditions and encompassing eight plankton size fractions ranging from 0.8 μm to 2 mm^36^ (Table S1). Notably, bacterial metagenome-assembled genomes (MAGs) characterized from this data set covered five distinct *Trichodesmium* populations^35^ (Figure 1, Table S2).

**Figure 1:**
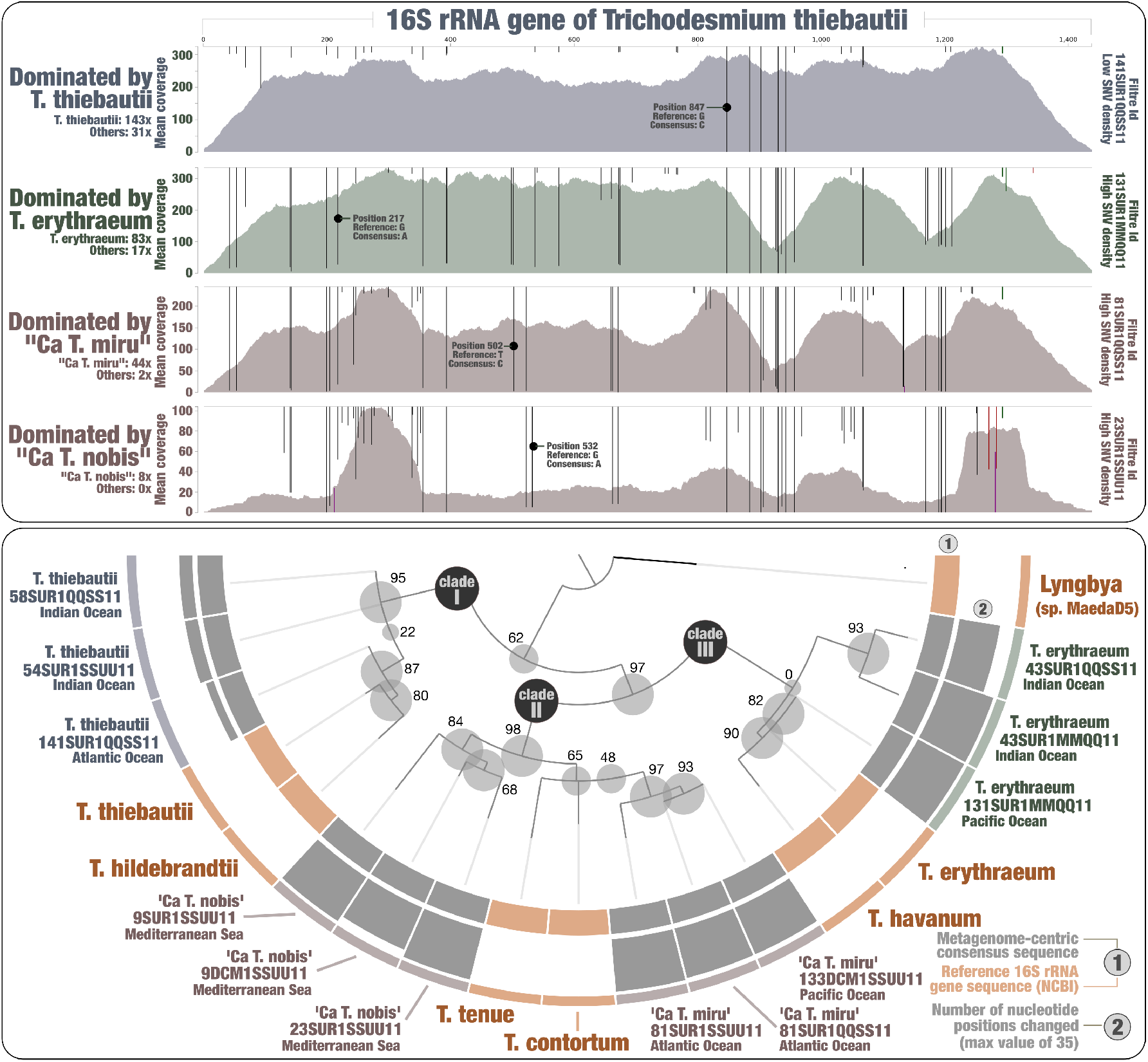
Hybrid phylogeny of the 16S rRNA gene of *Trichodesmium* species using references and metagenomes. Top panel displays read recruitments for the *T. thiebautii* 16S rRNA gene across four metagenomes, each dominated by a single *Trichodesmium* species (mapping stringency of >95% identity over >95% of the read length). Single nucleotide variants identified by anvi’o (when using default parameters) were visualized with an amplitude reaching 100% when all reads contained the same nucleotide type. Bottom panel displays a hybrid phylogenetic analysis using both reference sequences and 12 metagenome-centric consensus sequences. The number of changed nucleotide positions compared to the reference (*T. thiebautii* 16S rRNA gene) is presented for each consensus sequence. The phylogenetic tree was decorated with associated data and visualized using anvi’o.

The *Trichodesmium* MAGs were affiliated to *T. erythraeum* (one population), *T. thiebautii* (two closely related populations), and two candidate species we tentatively named ‘Ca. *Trichodesmium miru*’ (one population) and ‘Ca. *Trichodesmium nobis*’ (one population) that form a distinct evolutionary clade (ANI of 94%) distantly related from both *T. thiebautii* and *T. erythraeum* (ANI <91%) (Figure S1, Table S2). The five MAGs have a length ranging between 5.4 Mbp and 6.8 Mbp, with an estimated completion >90% (average of 96%). They were mostly detected in the Indian Ocean, with the exception of ‘Ca. *Trichodesmium miru*’ that occurred mostly in the Atlantic Ocean (Table S3). Furthermore, their niche partitioning between the Pacific Ocean, Red Sea and Mediterranean Sea revealed different distribution patterns between the four species. For instance, ‘Ca. *Trichodesmium nobis*’ was markedly more detected in the Mediterranean Sea. As expected, signal across size fractions indicates that all five populations mostly occur under the form of filaments and colonies in the sunlit ocean. We note that ‘Ca. *Trichodesmium miru*’ was substantially more detected in the largest size fraction (180-2000 μm), suggesting it forms larger aggregates in the Atlantic Ocean compared to the other *Trichodesmium* populations mostly detected in the Indian Ocean. In terms of overall signal across metagenomes regardless of oceanic regions or size fractions, the two candidate species were abundant but substantially less so than compared to *T. thiebautii* and *T. erythraeum* populations (Table S3). On the other hand, ‘Ca. *Trichodesmium miru*’ was detected in 29 *Tara* Ocean stations, being markedly more widespread compare to MAGs from the other species, which were detected in a number of stations ranging from 19 to 23 (Tables S2 and S3).

### Computing the metagenome-centric consensus of 16S rRNA genes

16S rRNA gene sequences are missing in most MAGs (including within *Trichodesmium*) due to their high evolutionary stability and occurrence in multi-copy (e.g.,^34^). While in uncharted territory, it is in theory possible to retrieve the missing gene of a bacterial population using a closely related reference to recruit metagenomic reads, provided this population dominates the signal within the range of the affiliated genus in the sequence space. Since each of the four *Trichodesmium* species dominated the metagenomic signal at times within the scope of this genus (e.g., in parts of the Atlantic Ocean for ‘Ca. *Trichodesmium miru*’), we took this opportunity to recover their 16S rRNA genes. We used the 16S rRNA gene of *T. thiebautii* as bait and performed a stringent read recruitment (to minimize non-specific mapping) for metagenomic triplicates targeting each species (Table S4). For each metagenome, gene positions that differed from the reference among recruited reads were changed accordingly to the most prevalent nucleotide type. Using this approach, we could create 12 metagenome-centric consensus 16S rRNA gene sequences for *Trichodesmium*. As expected, metagenomes dominated by *T. thiebautii* were highly coherent with the reference sequence, requiring only 5 to 8 changes that denote slight differences between the culture and dominant population in one hypervariable region (Figure 1). In contrast, metagenomes dominated by other species required between 25 and 35 changes covering multiple regions of the gene (Table S4), denoting this time greater distances between *Trichodesmium* species.

We then performed a phylogenetic analysis using the 12 consensus sequences along with reference 16S rRNA genes retrieved from NCBI, recapitulating the three *Trichodesmium* clades while nesting the candidate species into clade II. NCBI blast confirmed that the candidate species contain 16S rRNA gene signal most closely related to *T. tenue* and *T. contortum* among the metagenomes considered (percent identity >99% for ‘Ca. *Trichodesmium miru*’ and >98.5% for ‘Ca. *Trichodesmium nobis*’). Yet, we detected nucleotide differences in multiple regions of the 16S rRNA gene when using *T. tenue* and *T. contortum* as baits for mapping (Figure S2), suggesting that ‘Ca. *Trichodesmium miru*’ and ‘Ca. *Trichodesmium nobis*’ correspond to previously uncharacterized species within clade II of this genus.

### The case for Trichodesmium species lacking nitrogen fixation gene apparatus

We performed a comparative genomic survey of the genus *Trichodesmium* by considering the five newly identified MAGs plus two reference genomes from cultivation and accessed from NCBI: *T. erythraeum* IMS101 (closed genome) and *T. thiebautii* H9-4 (fragmented and only 72% complete). Our pangenomic analysis of these seven genomes with a total of 33,249 genes resulted in 7,778 gene clusters (Figure 1, Table S5). We collapsed singletons (2,542 gene clusters only detected in a single genome) and grouped some of the remaining gene clusters into bins based on their occurrence across genomes: (1) a core-genome (2,183 gene clusters), (2) gene clusters characteristic of *T. erythraeum* and *T. thiebautii* (“*erythraeum/thiebautii*” bin; n=99), (3) and gene clusters characteristic of the two candidate species (“*miru/nobis*” bin; n=157). The overall pangenomic trends for these four *Trichodesmium* species revealed a relatively large pan-genome but also denoted species-specific gene clusters that might be linked to different life styles for *Trichodesmium* clades I, II and III.

In order to provide a global view of functional capabilities across four *Trichodesmium* species, we accessed functions in their gene content using Pfam^37^ within the anvi’o pangenomic workflow^38^ (Table S5), COG20 functions, categories and pathways^39^, KOfam^40^, KEGG modules and classes^41^ within the anvi’o genomic workflow^42^ (Table S6), and RAST annotation^43^ (Table S7). The two most prominent functional capabilities of *Trichodesmium* are photosynthesis and nitrogen fixation. As expected, a large set of photosynthetic genes occurred in the seven genomes. On the other hand, multiple lines of evidence - emerging from the inspection of all functional annotations and cross-validated by complementary metagenomic investigations - point to ‘Ca. *Trichodesmium nobis*’ and ‘Ca. *Trichodesmium miru*’ lacking the ability to fix nitrogen from the atmosphere, providing a first case for the occurrence, in plain sight, of non-diazotrophic marine *Trichodesmium* species.

First and foremost, MAGs corresponding to the two candidate species lacked the entire nitrogen fixation gene apparatus, with the corresponding gene clusters occurring in the “*erythraeum/thiebautii*” pangenomic bin (Figure 2). This lack of signal was recapitulated across all functional annotations (Table S8) after removing few false positives easily identified by NCBI blast (Table S6). For instance, we found that COG20 functions incorrectly annotated as “Nitrogenase ATPase subunit NifH / coenzyme F430 biosynthesis subunit CfbC” a gene corresponding in reality to “ferredoxin : protochlorophyllide reductase”. Just two *nifU* related genes were detected (with NCBI blast confirmation) in those MAGs, however their occurrence in non-diazotrophic lineages indicates they are not reliable markers for nitrogen fixation. More relevant nitrogen fixation gene markers (*nifHDK and nifENB*) were successfully detected in a wide range of newly identified marine diazotrophic MAGs spanning multiple phyla^34,35^, suggesting our functional workflow could detect those markers in newly identified *Trichodesmium* species. It did not.

**Figure 2:**
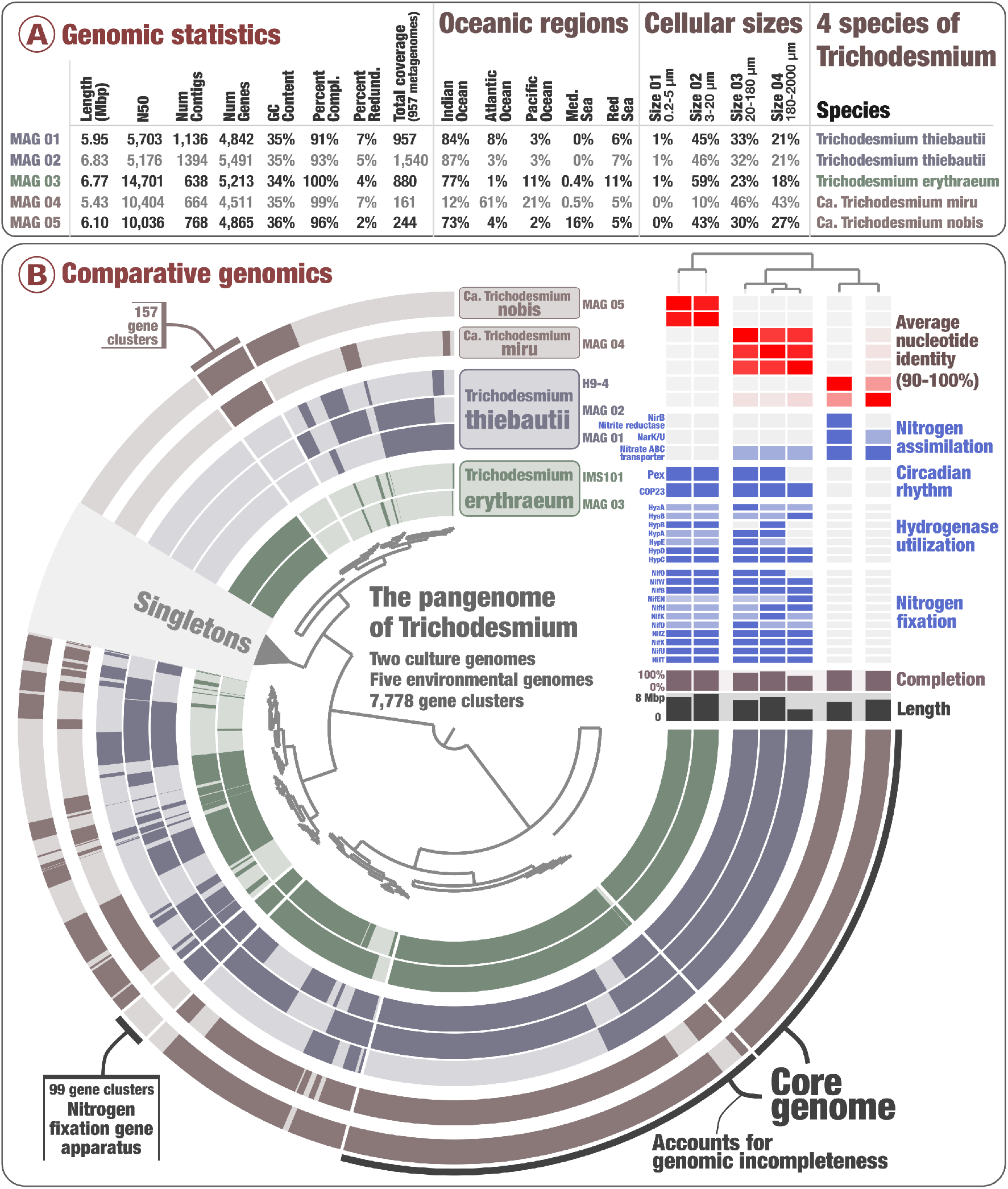
The pangenome of *Trichodesmium*. Panel A displays genomic statistics for the five environmental *Trichodesmium* MAGs, along with their environmental signal among 937 *Tara* Oceans metagenomes. Panel B displays the *Trichodesmium* pangenome that covers 33,249 genes and 7,776 gene clusters from seven genomes of *Trichodesmium* corresponding to four distinct species. The top-right corner of panel B holds average nucleotide identity between genomes and a selection of RAST functional features decorate this pangenome, which was visualized with the anvi’o interactive interface^42^. Finally, genomes were organized based on the average nucleotide identity metric.

Perhaps more problematic, reconstructing genomes from metagenomes often suffers from quality and completion issues, which could explain the lack of *nif* genes in some *Trichodesmium* MAGs. As a first effort to address this, we investigated the occurrence of a comprehensive marine *nifH* gene database (see methods) across the *Tara* Oceans metagenomes, which includes *nifH* genes for T. *erythraeum* and *T. thiebautii* with >95% sequence identity. We used a mapping stringency sufficiently low (>80% sequence identity over 80% of the read length) to capture this genus in the nucleotidic sequence space. The *nifH* gene sequence has long been documented to display very low genetic diversity among *Trichodesmium* species^18,44,45^. Yet, we found that *Tara* Oceans metagenomes with high signal for ‘Ca *Trichodesmium miru*’ did not contain the expected signal for *Trichodesmium nifH* genes (Figure 3 and Table S3). For instance, at Station 133 in the Pacific Ocean (deep chlorophyll maximum layer, 180-2000 μm size fraction) where only ‘Ca *Trichodesmium miru*’ was detected with a mean coverage of 13.8X (i.e., its genome was sequenced nearly 14 times in this metagenome), not a single metagenomic read matched to the >750 nucleotides long *nifH* genes from *T. erythraeum* and *T. thiebautii*, or any of the other *nifH* genes in the database. Overall, the correlation between the coverages of *Trichodesmium* genomes and their *nifH* genes better correlated in those samples when excluding the two candidate species (R^2^:0.99). ‘Ca *Trichodesmium nobis*’ displayed a higher niche overlap with *T. erythraeum* and *T. thiebautii*, nevertheless we could detect a clear discrepancy between its genomic occurrence and signal for *Trichodesmium nifH* genes in metagenomes of the Mediterranean Sea (Figure 3, Table S3). To expand this search for missing signal beyond the *nifH* gene, we then mapped metagenomic reads against the entire closed genome of *T. erythraeum* using an even lower mapping stringency (70% identity over 70% of the read length), revealing this time a lack of signal for the entire *nif* operon in samples dominated by the candidate species (Figure 4). In contrast, in samples that were dominated by *T. thiebautii*, the entire nif operon was perfectly recovered, except for the divergent intergenic regions. Thus, metagenomic signal for *nifH* and related genes were coherent with the comparative genomic insights, both pointing to a lack of the nitrogen fixation gene apparatus in the two closely related candidate species.

**Figure 3:**
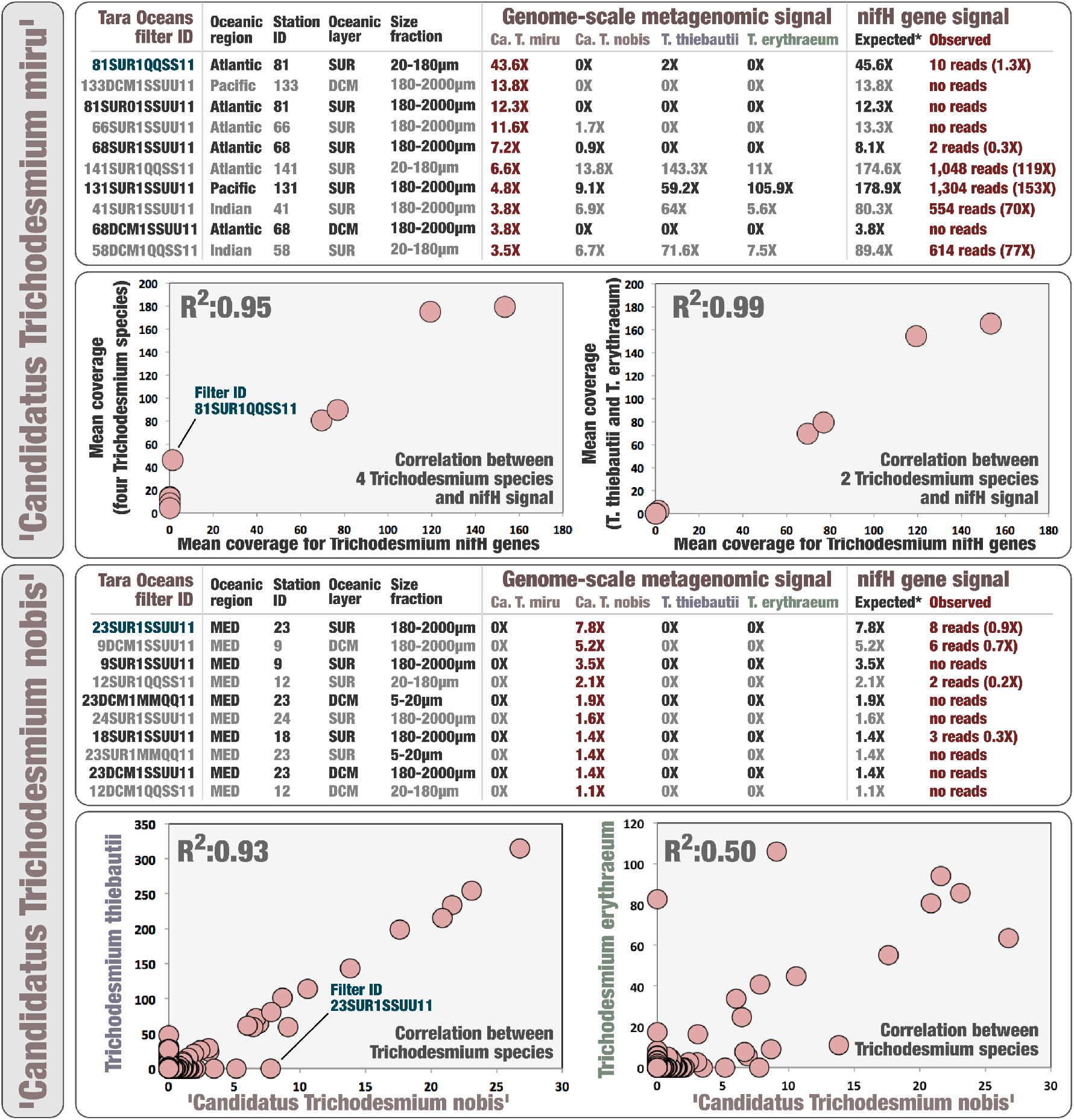
Metagenomic signal for *nifH* gene in the context of ‘Ca *Trichodesmium miru*’ and ‘Ca *Trichodesmium nobis*’. Top panel summarizes metagenomic read recruitment results for five *Trichodesmium* MAGs (genome-scale with mapping stringency of >90% identity over >80% of the read length) and three known *Trichodesmium nifH* genes (gene-centric with mapping stringency of >80% identity over >80% of the read length) across ten *Tara* Oceans metagenomes with highest mean coverage for ‘Ca *Trichodesmium miru*’. It also shows the correlation between mapped metagenomic reads for Trichodesmium *nifH* genes and either the cumulative mean coverage of all four Trichodesmium species (five MAGs) or the cumulative mean coverage of just *T. thiebautii* and *T. erythraeum* (three MAGs) across the same set of ten metagenomes. Bottom panel displays the correlation between ‘Ca *Trichodesmium nobis*’ and two *Trichodesmium* species across all *Tara* Oceans metagenomes. It also summarizes metagenomic read recruitment results for five *Trichodesmium* MAGs (genome-scale) and three known *Trichodesmium nifH* genes (gene-centric) across ten *Tara* Oceans metagenomes with highest mean coverage for ‘Ca *Trichodesmium miru*’ and no detection of the other MAGs (MED: Mediterranean Sea).

**Figure 4:**
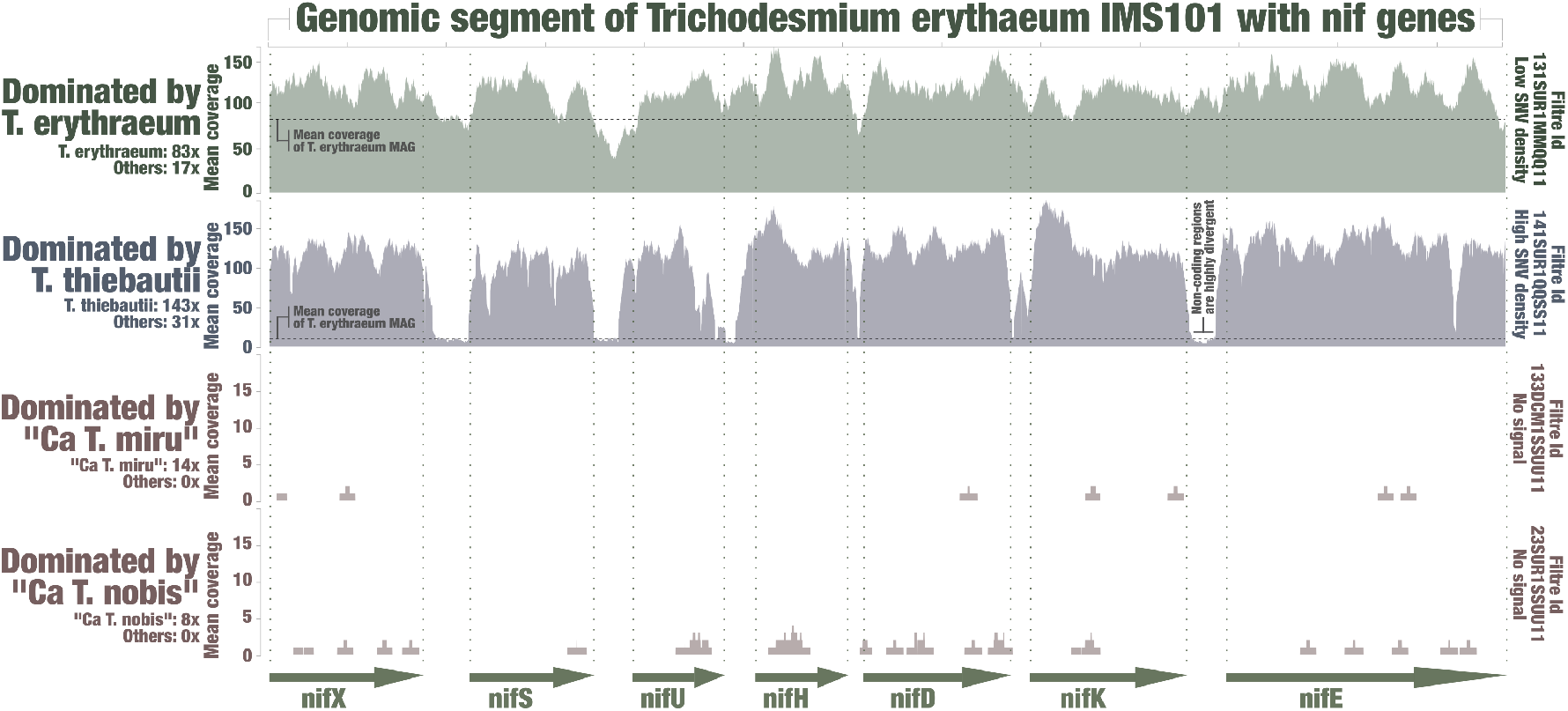
Metagenomic signal for the nif operon of *Trichodesmium erythraeum*. The figure displays read recruitment for the *T. erythraeum* IMS101 nif operon across four metagenomes, each dominated by a single *Trichodesmium* species (low mapping stringency of >70% identity over >70% of the read length; anvi’o visualization).

Hydrogen is a by-product of nitrogen fixation that cyanobacterial diazotrophs reutilize to gain energy^28–30^. We found that functions related to the recycling of hydrogen (*hyaABD* and *hypABCDE*,) were only missing in MAGs corresponding to the two candidate species (Figure 2B, top right panel; Table S8), which was also supported by a lack of metagenomic read recruitments for the *hyaABD* and *hypABCDE* genes of *T. erythraeum* in relevant metagenomes with low mapping stringency, echoing results for the *nif* operon (Figure S3). Non-diazotrophs in the surface ocean may have no reasons to contain genes for the recycling of hydrogen, however they do need to assimilate biologically available nitrogen molecules. We found the nitrite/nitrate transporter gene *nark* in single copy in *T. erythraeum* and *T. thiebautii* (COG20 functions), yet it occurred in two copies in ‘Ca *Trichodesmium miru*’ and in three copies in ‘Ca *Trichodesmium nobis*’. This function provides a means to catalyze the uptake of nitrogen-enriched small molecules^46^. RAST annotation also detected an enrichment of this functional annotation in the candidate species. Furthermore, COG20 functions and RAST annotation identified the gene *nirB* allowing dissimilatory nitrate reduction (anaerobic respiration, nitrogen metabolism) in ‘Ca *Trichodesmium miru*’. NCBI best blast hits linked this gene to the distantly related phyla Lentisphaera and Verrucomicrobia (including *Rubritalea marina* isolated from a sponge^47^), rather than more closely related lineages within Cyanobacteria. In addition, the gene appears next to a cyanobacterial transposase. Thus, we observed a lack of hydrogen recycling genes concomitant with the enrichment of genes related to nitrogen assimilation and metabolism in the candidate species, coherent with a non-diazotrophic life style.

Some cyanobacterial diazotrophs possess circadian rhythms that regulate nitrogen fixation and photosynthesis activities temporally and spacially^48–51^. While all four *Trichodesmium* species contained genes coding for the core circadian clock proteins (*kaiABC*) linked to photosynthesis regulation^51^, the pangenomic analysis revealed two gene clusters corresponding to the circadian oscillating protein *COP23* and only missing in MAGs corresponding to the candidate species (Table S8). Furthermore, RAST annotation only identified one gene for *COP23* and one gene for Pex (another function linked to circadian rhythm), which were only detected in *T. erythraeum* and *T. thiebautii* (Figure 2B, top right panel). The cyanobacterial nitrogen fixation rhythm can be disconnected from light variations in laboratory experiments^23,52^, however genes related to this mechanism have yet to be fully understood. Our results echo previous work on the *COP23* and *Pex* genes^53–56^, suggesting that at least some of these genes play a central role in the circadian rhythm of nitrogen fixation in *T. erythraeum* and *T. thiebautii*.

Finally, we cross-validated these insights by performing six additional *Tara* Oceans genome-resolved metagenomic surveys guided by the known distribution of four *Trichodesmium* species and designed to only target either ‘Ca *Trichodesmium miru*’ or ‘Ca *Trichodesmium nobis*’ by means of single assemblies or small co-assemblies (Table S9). First, we used HMM models for the *nifHDK* and *nifENB* genes designed to cover the entire bacterial spectrum and found no trace of these gene markers in the six raw metagenomic assemblies, which contain contigs as short as 1,000nt (as a perspective, the *nifB* gene of *T. erythraeum* is 1,470 nt long). Then, we characterized and manually curated the *Trichodesmium* MAG of the targeted candidate species in each metagenomic assembly. Their completion ranged between 90.1% and 98.6%, with N50 as high as 26,010nt (Table S9). Their genetic content agreed with the previous lines of evidence as they were enriched in genes for nitrate/nitrite transporter *nark*, contained this time the gene *nirB* in both species (systematically linking this gene to Lentisphaera and Verrucomicrobia), and most importantly lacked genes related to the nitrogen fixation apparatus (except for the *nifU* related genes) and hydrogen recycling (*hyaABD* and *hypABCDE*) (Table S10). Thus, our lines of evidence for non-diazotrophic *Trichodesmium* species were reproducible using various metagenomic combinations and are supported by a lack of signal for *nifHDK* and *nifENB* genes in the raw metagenomic assemblies.

## Discussion

The association between *Trichodesmium* and nitrogen fixation has been deeply rooted in our minds due to considerable culture and fieldwork legacies, and as a result *Trichodesmium* is routinely referred to as a diazotrophic genus (e.g.,^14^). Here, we provide multiple lines of evidence indicating that the newly identified species ‘Ca. *Trichodesmium miru*’ and ‘Ca. *Trichodesmium nobis*’, characterized by means of genome-resolved metagenomics and found abundant in multiple regions of the open ocean, do not have the ability to fix nitrogen from the atmosphere. These species form a sister genomic clade most closely related to *T. tenue* and *T. contortum* based on metagenome-centric 16S rRNA gene consensus sequences (Figure 1). Near-complete environmental genomes for those candidate species denoted a lack of the entire nitrogen fixation gene apparatus and hydrogen recycling genes concomitant with the enrichment of genes related to nitrogen assimilation. These comparative genomic insights were supported by the absence of metagenomic signal for *Trichodesmium nifH* genes, the reference genome *T. erythraeum* IMS101, and targeted metagenomic assemblies and binning efforts, contrasting with the current paradigm that *Trichodesmium* species are necessarily capable of nitrogen fixation.

When considering ‘Ca. *Trichodesmium miru*’ and ‘Ca. *Trichodesmium nobis*’ as non-diazotrophic species, their detection mostly restricted to large size fractions echoes trends for known *Trichodesmium* species and suggests filaments and colonies are the norm for this genus regardless of nitrogen fixation, albeit this has yet to be demonstrated. But most importantly, a critical consequence of this gene loss is that *Trichodesmium* might have a broader marine niche distribution compared to its nitrogen fixation capability. In fact, *T. erythraeum* and *T. thiebautii* remained undetected in 27 stations with clear signal for the putative non-diazotrophic *Trichodesmium* species (based on genome-wide metagenomic read recruitment, see Figures 5 and S4, Table S3). These stations cover the Indian, Atlantic and Pacific Oceans as well as the Mediterranean Sea, which indicates that ‘Ca. *Trichodesmium miru*’ and ‘Ca. *Trichodesmium nobis*’ are not only widespread but might also substantially expand the ecological niche of this genus. In our view, this stresses the need to disconnect taxonomic signal for *Trichodesmium* from the ability of a microbial community to fix nitrogen. Far from contesting the paramount importance of *T. erythraeum* and *T. thiebautii* for nitrogen fixation in the open ocean (in the context of dozens of other abundant cyanobacterial and heterotopic bacterial populations^35^), we merely suggest that differentiating diazotrophic from non-diazotrophic *Trichodesmium* populations might be needed to refine our understanding of the nitrogen balance in the oceans and seas. Counting *Trichodesmium* cells to survey the biomass of marine diazotrophs, as routinely performed for decades (e.g.,^57^), might lead to erroneous nitrogen fixation rate estimations, provided of course that non-diazotrophic *Trichodesmium* species indeed do exist and are not just a metagenomic mirage. In addition, the joint study of diazotrophic and non-diazotrophic *Trichodesmium* populations under culture conditions could bolster our understanding of the underlying genetic mechanisms for nitrogen fixation in *Trichodesmium* and beyond. As a humble step in this direction, it could be hypothesized that some circadian rhythm genes missing in ‘Ca. *Trichodesmium miru*’ and ‘Ca. *Trichodesmium nobis*’ but present in *T. erythraeum* and *T. thiebautii* are specifically linked to the regulation of nitrogen fixation, rather than photosynthesis, among *Trichodesmium* diazotrophs.

**Figure 5:**
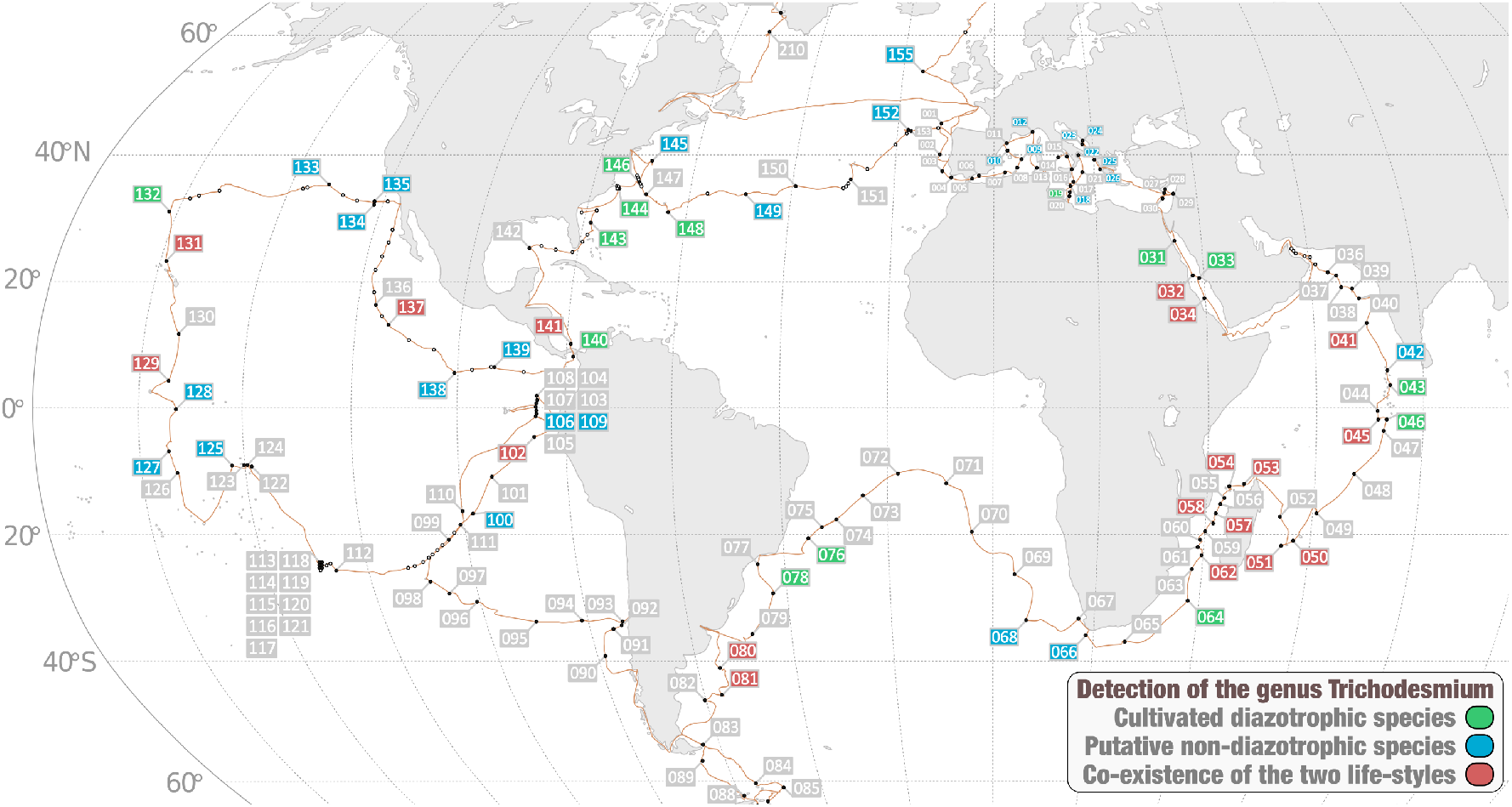
Detection of *Trichodesmium* species across the *Tara* Oceans stations. The world map describes stations in which we detected (1) only *T. erythraeum* and/or *T. thiebautii* (diazotrophic species), (2) only ‘Ca. *Trichodesmium miru*’ and/or ‘Ca. *Trichodesmium nobis*’ (putative non-diazotrophic species), (3) or species from both groups. This surveys covers all size fractions >0.2 μm.

Non-diazotrophic *Trichodesmium* populations might have diverged from a diazotrophic lineage in order to fill distinct ecological niches at the surface of the open ocean (gene loss hypothesis), echoing within a genus results observed at the level of Cyanobacteria that revealed its complex evolutionary history^14,58,59^ and suggested repeated losses of the *nif* genes within this phylum^60^. Interestingly, ‘Ca. *Trichodesmium nobis*’ strongly correlated with *T. thiebautii* (R^2^=0.93), the most abundant of the four species within the scope of *Tara* Oceans metagenomes. The considerable overlap between these two species in the Indian Ocean especially, with one prevalent diazotroph and a less abundant one (ratio of about 1/15) lacking the ability to fix nitrogen fits well with the principles behind the black queen hypothesis^61^. This hypothesis states that gene loss can provide a selective advantage as long as the function is dispensable. ‘Ca. *Trichodesmium nobis*’ might have lost its ability to fix nitrogen in part because it could benefit from the nitrogen fixation of *T. thiebautii,* the “leaky helper” mentioned by Morris et al.^61^. Furthermore, the *nirB* gene linked to dissimilatory nitrate reduction in low-oxygen environments might provide a clue to understanding mechanisms behind this ecologically important evolutionary process, which based on our lines of evidence occurred in some *Trichodesmium* species but not in others. First, this gene was only detected in the two candidate species and is most closely related to Lentisphaera and Verrucomicrobia related genes. Second, *nirB* genes have been observed in the genomic content of non-cyanobacterial epibionts living at the surface of *Trichodesmium* colonies^62,63^, suggesting a possible complementary functional role contributing to their interactions. The temporality of *nirB* gene acquisition and loss of nitrogen fixation and hydrogen recycling genes in the candidate species are currently unknown. Nevertheless, one could wonder if lateral gene transfers between *Trichodesmium* colonies and their epibionts have provided a non-diazotrophic evolutionary path for the common ancestor of ‘Ca *Trichodesmium miru*’ and ‘Ca *Trichodesmium nobis*’, opening exciting prospects regarding the ecology and evolution of this model planktonic lineage in the context of gene flows.

## Conclusion

The cyanobacterial genus *Trichodesmium* includes some of the most prominent marine nitrogen fixing species, which have been extensively studied in culture and the field for decades. Here, we explored the genome-resolved metagenomic content of two previously uncharacterized *Trichodesmium* species relatively abundant in the surface of oceans and seas. Critically, multiple lines of evidence point to the existence of non-diazotrophic *Trichodesmium* species with distinct ecological niches. Establishing that newly identified microbial populations lack critical functional traits from the sole perspective of environmental genomics could be perceived as a precarious endeavor. This is especially true when observations are not in line with those resulting from decades of cultivation. Yet, this approach comes with its own strengths, including reference-free de novo metagenomic assemblies on one hand and metagenomic read recruitments using reference genomes and gene markers as bait on the other. In the case of ‘Ca. *Trichodesmium miru*’ and ‘Ca. *Trichodesmium nobis*’, genome-resolved metagenomics and read recruitments applied to the considerable metagenomic legacy of *Tara* Oceans provided strong evidence that two closely related *Trichodesmium* species have lost the ability to fix nitrogen from the atmosphere. Culture representatives for these candidate species are needed to confront and move beyond those metagenomic insights. Until then, we are reminded that some long-established links between taxonomic lineages and functional traits supporting our understanding of the ocean microbiome might not always hold true.

## Material and methods

### *Tara* Oceans metagenomes

We analyzed a total of 937 *Tara Oceans* metagenomes available at the EBI under project PRJEB402 (https://www.ebi.ac.uk/ena/browser/view/PRJEB402). Table S1 reports general information (including the number of reads and environmental metadata) for each metagenome.

### Biogeography of MAGs

We performed a mapping of all metagenomes to calculate the mean coverage and detection of MAGs. Briefly, we used BWA v0.7.15 (minimum identity of >90% over >80% of the read length) and a FASTA file containing the 1,888 non-redundant MAGs from Delmont et al.^35^ to recruit short reads from all 937 metagenomes. We considered MAGs were detected in a given filter when >25% of their length was covered by reads to minimize non-specific read recruitments^34^. The number of recruited reads below this cut-off was set to 0 before determining vertical coverage and percent of recruited reads.

### Metagenome-centric consensus 16S rRNA gene sequences

We performed a mapping of metagenomes against a reference 16S rRNA gene in order to retrieve corresponding signal for each of the four *Trichodesmium* species. Briefly, we used BWA v0.7.15 (minimum identity of >95% over >95% of the read length) and a FASTA file containing the 16S rRNA gene sequence T. *thiebautii* (AF013027) to recruit short reads from 12 metagenomes. We then used the anvi’o v.7 metagenomic workflow^42^ to create a CONTIG database (the reference 16S rRNA gene sequence) and PROFILE databases for each metagenome, with the flag “--report-variability-full” in order to report every single nucleotide variations (full mode), or without it (default mode). After merging the PROFILE databases, we used the anvi’o interactive interface with inspection mode to visualize the coverage of this gene across metagenomes in the context of single nucleotide variants (default mode). We then used the program “anvi-gen-gene-consensus-sequences” with the flag “--contigs-mode” to generate metagenome-centric consensus 16S rRNA gene sequences (full mode).

### Phylogenetic inferences using 16S rRNA gene sequences

We performed a phylogenetic analysis of the metagenome-centric consensus 16S rRNA gene sequences (see previous section) and reference 16S rRNA genes retrieved from NCBI. Briefly, we used the online platform GenomeNet (https://www.genome.jp/) to generate a phylogenetic tree at the nucleotide level and using as parameters the function "build" of ETE3 v3.1.1^64^, MAFFT v6.861b with the linsi options^65^ for alignment. Columns with more than ten percent of gaps were removed from the alignment using trimAl v1.4.rev6^66^. Finally, ML tree was inferred using PhyML v20160115 ran with model GTR and parameters “-o tlr --alpha e --bootstrap 100 --pinv e --nclasses 4 -f m”^67^. Branch supports were computed out of 100 bootstrapped trees. We used anvi’o to visualize the phylogenomic tree in the context of additional information.

### Phylogenomic analysis of cyanobacterial genomes

We used PhyloSift^68^ v1.0.1 with default parameters to infer associations between genomes in a phylogenomic context. Briefly, PhyloSift (1) identifies a set of 37 marker gene families in each genome, (2) concatenates the alignment of each marker gene family across genomes, and (3) computes a phylogenomic tree from the concatenated alignment using FastTree^69^ v2.1. We used anvi’o to visualize the phylogenomic tree in the context of additional information.

### Pangenomic analysis of MAGs

We used the anvi’o pangenomic workflow^38^ to compute and visualize the pangenome of *Trichodesmium*. Briefly, the workflow consisted of three main steps: (1) we generated an anvi’o genome database to store DNA and amino acid sequences, as well as functional annotations of each gene in the seven *Trichodesmium* genomes under consideration, (2) we computed the *Trichodesmium* pangenome from a genome database by identifying ‘gene clusters’, and (3) we displayed the pangenome to visualize the distribution of gene clusters across genomes. The gene clusters represent sequences of one or more predicted open reading frames grouped together based on their homology at the translated DNA sequence level. To compute the *Trichodesmium* pangenome, we used the program‘anvi-pan-genome’with the flag‘--use-ncbi-blast’and default parameters. This program (1) calculates similarities of each amino acid sequence in every genome against every other amino acid sequence using blastp^70^, (2) removes weak hits using the ‘minbit heuristic’, which was originally described in ITEP^71^, (3) uses the MCL algorithm^72^ to identify gene clusters in the remaining blastp search results, (4) computes the occurrence of gene clusters across genomes and the total number of genes they contain, (5) performs hierarchical clustering analyses for gene clusters (based on their distribution across genomes) and for genomes (based on gene clusters they share) using Euclidean distance and Ward clustering by default, and finally (6) generates an anvi’o pan database that stores all results for downstream analyses and was used to visualize the *Trichodesmium* pangenome in the interactive interface.

### Functional inferences of MAGs

We inferred functions among the *Trichodesmium* genes using (1) Pfam^37^ from within the anvi’o pangenomic workflow, (2) COG20 functions, categories and pathways^39^, KOfam^40^, KEGG modules and classes^41^ within the anvi’o genomic workflow^42^ and (3) the RAST online platform^43^. Regarding the KEGG modules, we calculated their level of completeness in each genomic database using the anvi’o program “anvi-estimate-metabolism” with default parameters. The URL https://merenlab.org/m/anvi-estimate-metabolism describes this program in more detail. Lastly, we used online NCBI blasts to identify false positives regarding specific nitrogen fixation gene markers. False positives correspond to functional annotations for which NCBI blast identified a different function with lower e-values and bit scores.

### Metagenomic signal for the extended nifH gene database

We performed a mapping of metagenomes to calculate the mapped reads and mean coverage of sequences in an extended *nifH* gene database (see^35^ for more details). Briefly, we used BWA v0.7.15 (minimum identity of 80%) and a FASTA file containing the sequences to recruit short reads from 937 *Tara* Oceans metagenomes.

### Metagenomic signal for the reference *Trichodesmium erythraeum* IMS101

We performed a mapping of metagenomes against the reference genome *Trichodesmium erythraeum* IMS101 in order to retrieve corresponding signal for each of the four *Trichodesmium* species. Briefly, we used BWA v0.7.15 (minimum identity of >70% over >70% of the read length) and a FASTA file containing the entire genome *Trichodesmium erythraeum* IMS101 to recruit short reads from 12 metagenomes. We then used the anvi’o metagenomic workflow to create a CONTIG database (the genome) and PROFILE databases for each metagenome. After merging the PROFILE databases, we used the anvi’o interactive interface with inspection mode to visualize the coverage of regions of interest (e.g., the nif operon) across metagenomes.

### Reproducing the recovery of MAGs for the candidate species

As a supplement of the initial large-scale genome-resolved metagenomic survey^35,36^, we also performed additional genome-resolved metagenomic surveys by taking advantage of our knowledge of their distribution across the *Tara* Oceans metagenomes. Briefly, we used metagenomic reads as inputs for six metagenomic single assemblies or co-assemblies using MEGAHIT^73^ v1.1.1, and simplified the scaffold header names in the resulting assembly outputs using anvi’o^42^. We then completed the anvi’o manual binning workflow to extract *Trichodesmium* MAGs from these assemblies. For each targeted species, we used a distinct set of 8 relevant metagenomes for mapping in order to effectively use differential coverage for binning. Finally, we studied the functional repertoire of recovered *Trichodesmium* MAGs using anvi’o, as described in the “Functional inferences of MAGs” section.

### Search for *nifHDK* and *nifENB* genes in raw assemblies

We used HMM models designed to cover the entire bacterial spectrum of *nifHDK* and *nifENB* genes to search for these gene markers in raw metagenomic assemblies performed to extract the genomic content of the candidate species. The e-value cut-off was set to e-100.

## Supplemental information

We provided a supplemental information document describing the anvi’o workflows used in the study. Each section of the document includes the list of anvi’o programs and a breif explanation of the workflow.

## Data availability

First, the genomic resource (bacterial and archaeal MAGs from the surface of the oceans) is publicly available at http://www.genoscope.cns.fr/tara/. The link provides access to the 11 raw metagenomic co-assemblies from Delmont et al.^36^ as well as the FASTA file for 1,888 MAGs, including the five corresponding to *Trichodesmium*. In addition, the doi https://doi.org/10.6084/m9.figshare.14207321 provides access to (0) FASTA files for the initial five *Trichodesmium* MAGs, (1) anvi’o files corresponding to the metagenomic read recruitments used to generate 16S rRNA metagenomic consensus sequences, (2) anvi’o CONTIGS databases for reference *Trichodesmium* MAGs and isolate genomes (including functional annotations), (3) anvi’o files corresponding to the *Trichodesmium* pangenome, (4) anvi’o files corresponding to the metagenomic read recruitments for reference genome *Trichodesmium* erythraeum IMS101 (different mapping stringencies included), (5) targeted genome-resolved metagenomic surveys for the candidate *Trichodesmium* species (contigs >2,5kbp, anvi’o files and summaries), (6) anvi’o CONTIGS databases for the *Trichodesmium* MAGs extracted from the targeted genome-resolved metagenomic surveys (including functional annotations), (7) HMM models for six nitrogen fixation gene markers, (8) the supplemental tables and information, (9) and finally the raw assemblies (contigs >1kbp) for targeted genome-resolved metagenomic surveys.

## Acknowledgments

I would like to thank Michael D. Lee, A. Murat Eren, Eric Pelletier and Jed A. Fuhrman for technical support, editorial suggestions and constructive discussions regarding *Trichodesmium* and nitrogen fixation in the sunlit ocean. I also would like to thank Paul Frémont regarding world map production for the relative distribution of *Trichodesmium* species. In addition, this survey was especially made possible by two scientific endeavors: the sampling and sequencing efforts by the *Tara* Oceans consortium, and the bioinformatics and visualization capabilities afforded by anvi’o. As a result, I would like to thank everyone that contributed to *Tara* Oceans and anvi’o over the years^74,75^. In addition, *Tara* Oceans (which includes the *Tara* Oceans and *Tara* Oceans Polar Circle expeditions) would not exist without the leadership of the *Tara* Oceans Foundation and the continuous support of 23 institutes (https://oceans.taraexpeditions.org/). Finally, part of the computation was performed using the platine, titane and curie HPC machine provided through GENCI grants (t2011076389, t2012076389, t2013036389, t2014036389, t2015036389 and t2016036389).

## Supplemental figure

**Figure S01:**
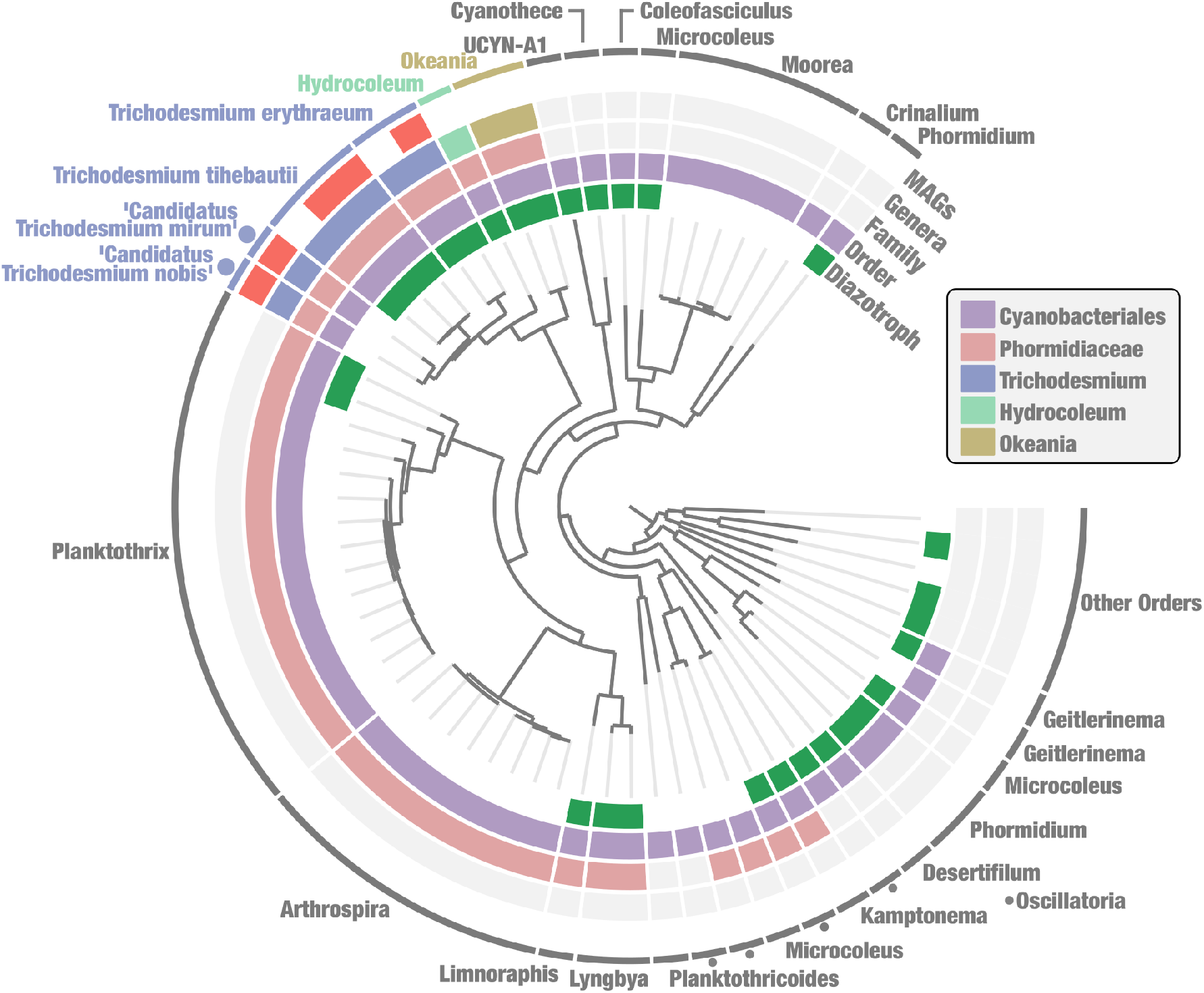
Phylogenomic analysis of cyanobacterial lineages closely related to *Trichodesmium*. This phylogenomic analysis includes xx reference genomes affiliated to Cyanobacteriales, including the five *Trichodesmium* population genomes characterized in this study (‘MAGs’ layer). The tree is rooted using five additional genomes affiliated to closely related orders within Cyanobacteria. Genomes containing nitrogen fixation genes are highlighted (‘diazotroph’ layer).

**Figure S02:**
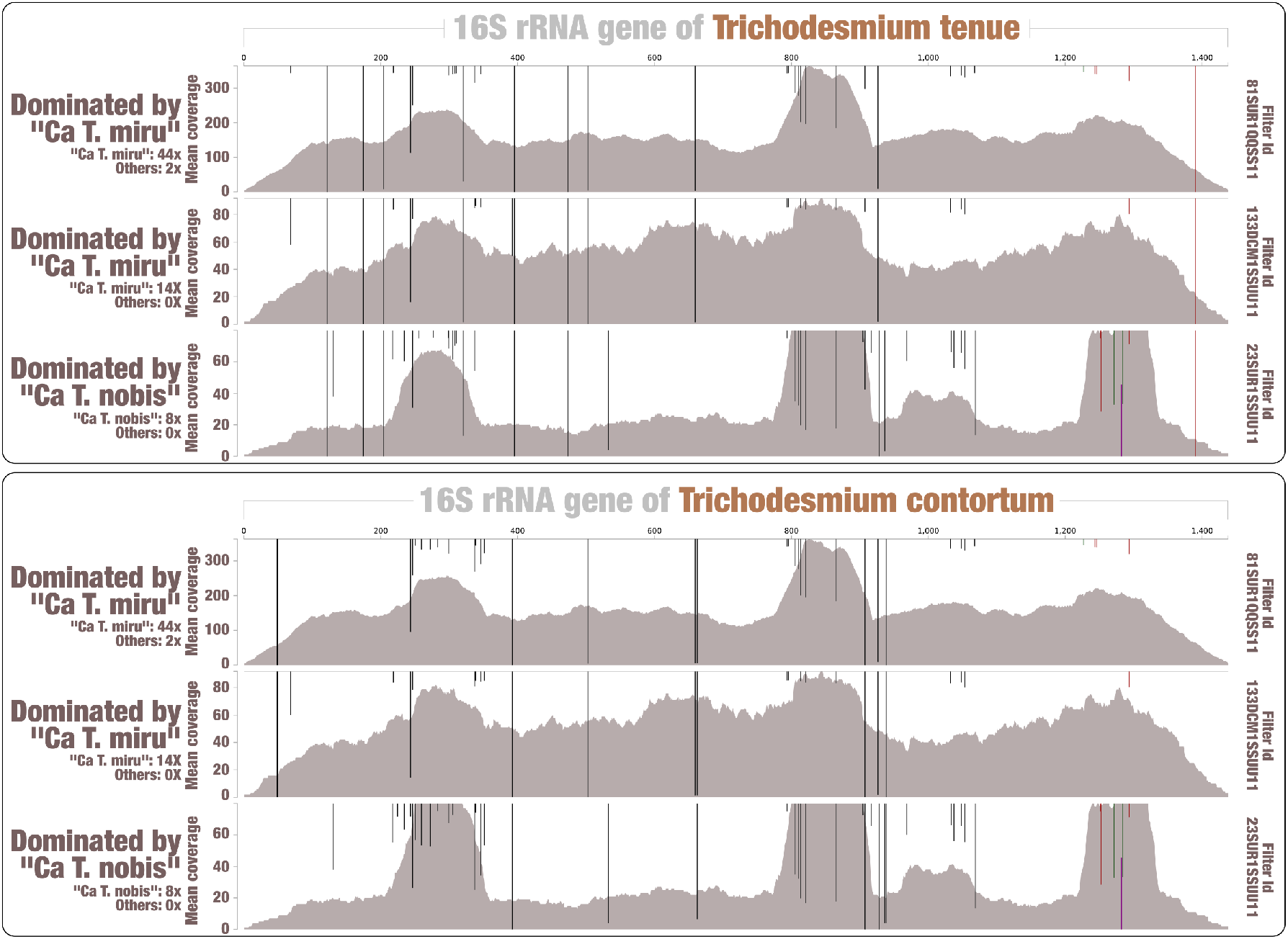
16S rRNA gene recruitment using Trichodesmium tenue and Trtichodesmium contortum as references. The figure displays read recruitment for the *T. tenue* and *T. contortum* 16S rRNA gene across three metagenomes dominated by candidate species (mapping stringency of >95% identity over >95% of the read length). Single nucleotide variants identified by anvi’o (when using default parameters) were visualized with an amplitude reaching 100% when all reads contained the same nucleotide type.

**Figure S03:**
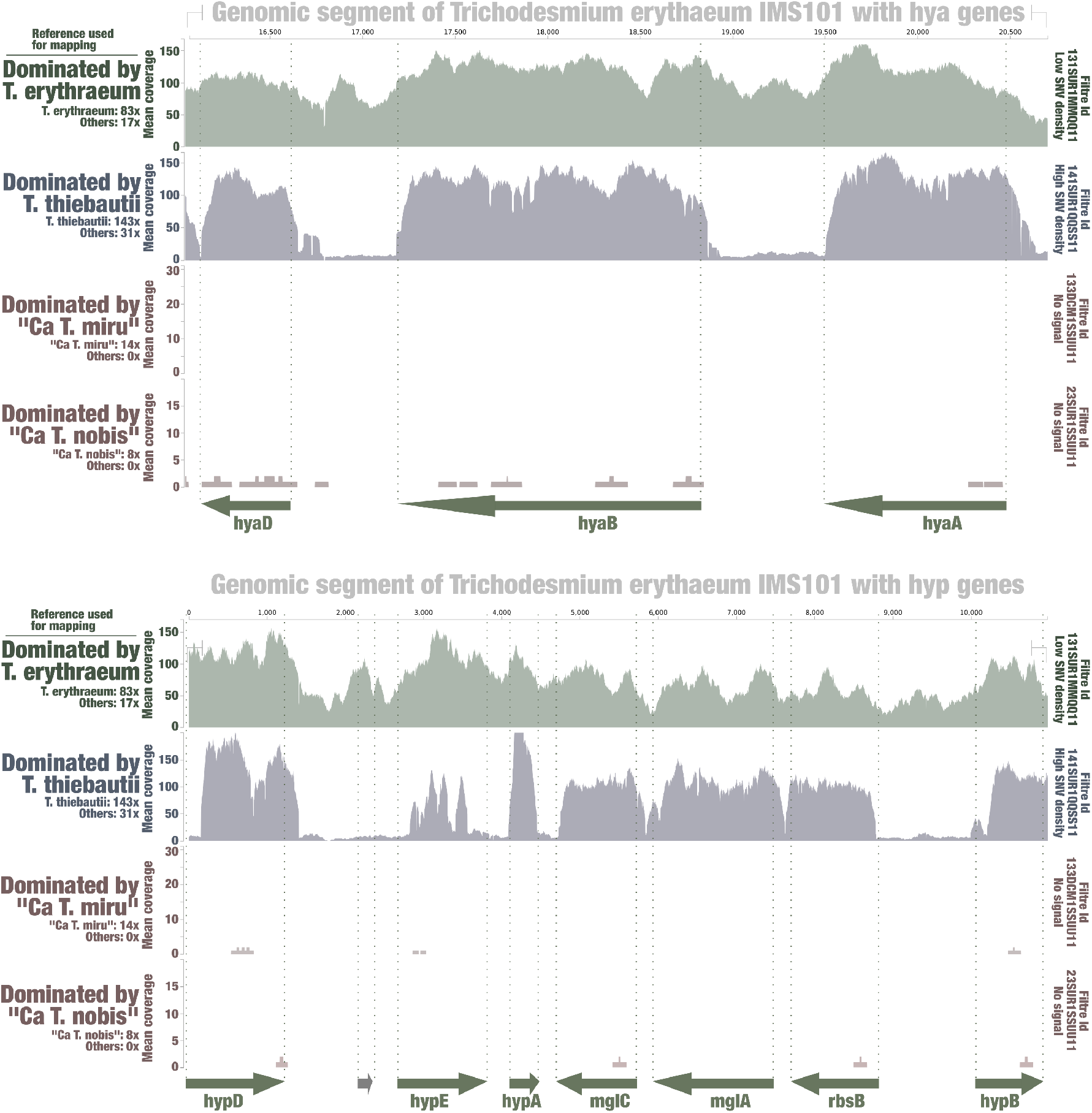
Metagenomic signal for the hya and hyp operons for hydrogen recycling of *Trichodesmium erythraeum*. The figure displays read recruitment for the *T. erythraeum* IMS101 hya and hyp operons across four metagenomes, each dominated by a single *Trichodesmium* species (low mapping stringency of >70% identity over >70% of the read length; anvi’o visualization).

**Figure S04:**
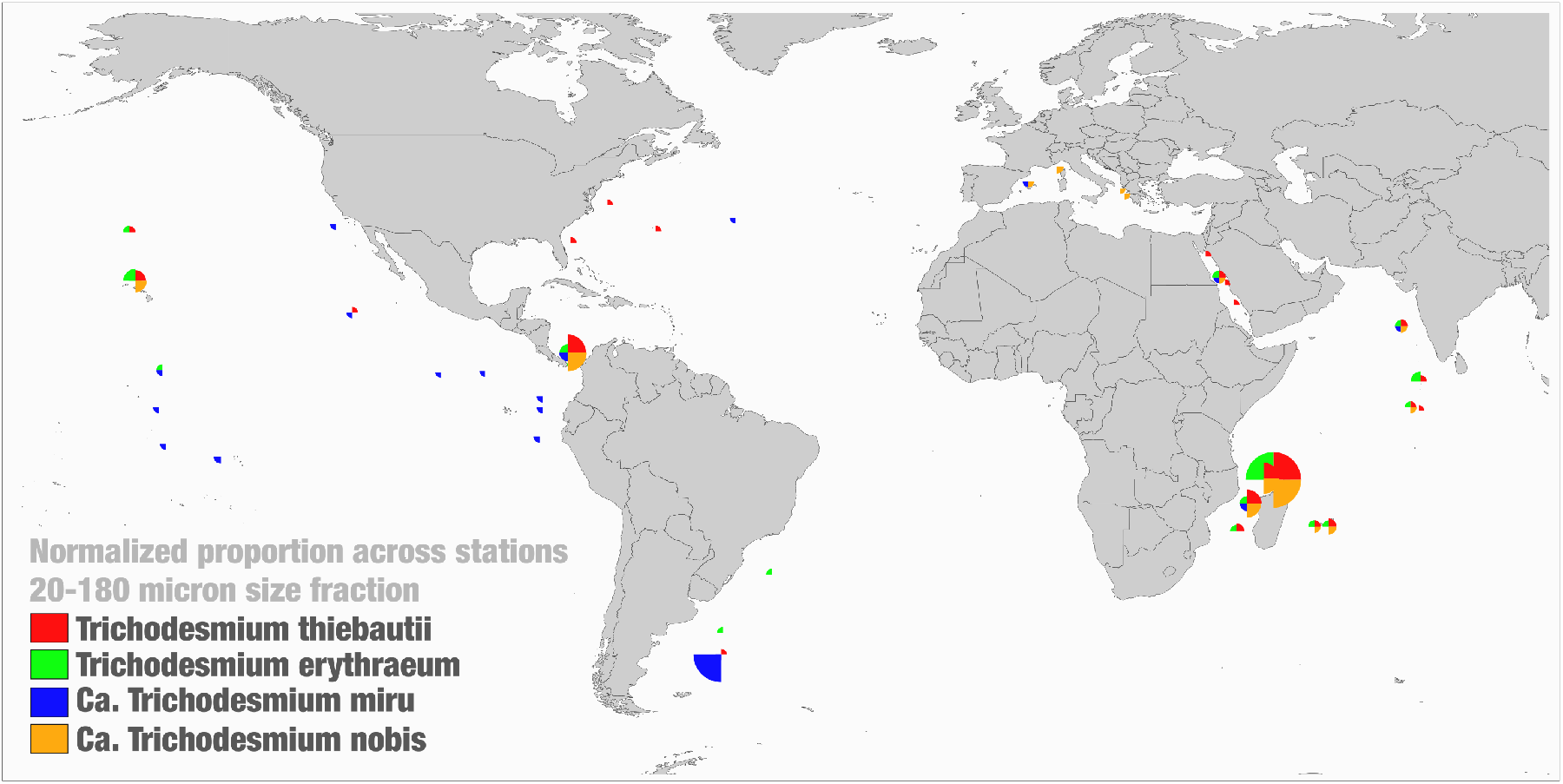
Normalized relative distribution of four *Trichodesmium* species. The world map displays the relative distribution of four *Trichodesmium* species across *Tara* Oceans metagenomes corresponding to the 20-180 microns size fraction (normalized so that each species has a full quarter of circle when it reaches its maximal value). The figure aims at providing some perspective regarding the occurrence of key marine *Trichodesmium* species in the surface of the open oceans and seas.

## Supplemental Tables

**Table S01:** Statistics for 937 *Tara* Oceans metagenomes organized by depth, size fraction and oceanic region. The table also contains environmental conditions across *Tara* Oceans stations.

**Table S02:** Statistics for the 1,888 bacterial and archaeal MAGs. The table contains genomic statistics (e.g., completion and length), taxonomic information, general mapping trends such as the cosmopolitan score, as well as additional information regarding the five *Trichodesmium* MAGs (e.g., ANI comparisons).

**Table S03:** Metagenomic read recruitment statistics for the five *Trichodesmium* MAGs (90% identity) and reference *nifH* genes (80% identity). The table contains mean genomic coverage values across the 937 *Tara* Oceans metagenomes.

**Table S04:** Metagenomic set used to generate the 12 metagenome-centric consensus sequences. The table describes the mean coverage of each *Trichodesmium* MAG and also contains reference and consensus 16S rRNA gene sequences.

**Table S05:** The pangenome of *Trichodesmium*. The table contains the anvi’o pangenomic summary outcome of seven *Trichodesmium* genomes (MAGs and isolates), connecting genes to gene clusters, genomes, bins and Pfam functional annotations.

**Table S06:** Functional annotation of seven *Trichodesmium* genomes (MAGs and isolates) using COG20 functions, categories and pathways, KOfam, KEGG modules and classes processed within anvi’o.

**Table S07:** Functional annotation of seven *Trichodesmium* genomes (MAGs and isolates) using the online RAST annotation platform.

**Table S08:** Summary of the lines of evidence across functional annotations for seven *Trichodesmium* genomes (MAGs and isolates) with respect to nitrogen fixation.

**Table S09:** Additional genome-resolved metagenomic surveys to recover MAGs from the two candidate *Trichodesmium* species. The table describes the sets of metagenomes used for assembly, along with the sets of metagenomes used for mapping.

**Table S10:** Functional annotation of redundant *Trichodesmium* MAGs for ‘Ca *Trichodesmium miru*’ (n=5) and ‘Ca *Trichodesmium nobis*’ (n=1) using using COG20 functions, categories and pathways, KOfam, KEGG modules and classes processed within anvi’o.

## Notes

https://www.genoscope.cns.fr/tara/

### Competing Interest Statement

The authors have declared no competing interest.

